# Demographic compensation does not rescue populations at a trailing range edge

**DOI:** 10.1101/117606

**Authors:** Seema Nayan Sheth, Amy Lauren Angert

## Abstract

As climate change shifts species' climatic envelopes across the landscape, equilibrium between geographic ranges and niches is likely diminishing due to time lags in demography and dispersal. If a species' range and niche are out of equilibrium, then population performance should decrease from cool, “leading” range edges, where populations are expanding into recently ameliorated habitats, to warm, “trailing” range edges, where populations are contracting from newly unsuitable areas. Population contraction signals that compensatory changes in vital rates are insufficient to buffer population growth from deteriorating environments. Life history theory predicts tradeoffs between fast development, high reproduction, and short longevity at low latitudes and slow development, less frequent but multiple bouts of reproduction, and long lifespan at high latitudes. If demographic compensation is driven by life history evolution, compensatory negative correlations in vital rates may be associated with this fast-slow continuum. An outstanding question is whether range limits and range contractions reflect inadequate compensatory life history shifts along environmental gradients, causing population growth rates to fall below replacement levels at range edges. We surveyed demography of 32 populations of the scarlet monkeyflower (*Erythranthe cardinalis*) spanning 11° latitude in western North America and used integral projection models to infer population dynamics and assess demographic compensation. Population growth rates decreased from north to south, consistent with leading-trailing dynamics. Southern populations are declining due to reduced survival, growth, and recruitment, despite compensatory increases in reproduction and faster life history characteristics, suggesting that demographic compensation will not rescue populations at the trailing range edge.

**SIGNIFICANCE STATEMENT:** While climate change is causing poleward shifts in many species' geographic distributions, some species' ranges have remained stable, particularly at low-latitude limits. One explanation for why some species' ranges have not shifted is demographic compensation, whereby declines in some demographic processes are offset by increases in others, potentially buffering populations from extinction. However, we have limited understanding of whether demographic compensation can prevent collapse of populations facing climate change. We examined the demography of natural populations of a perennial herb spanning a broad latitudinal gradient. Despite increases in reproduction, low-latitude populations declined due to diminished survival, growth, and recruitment. Thus, demographic compensation may not be sufficient to rescue low-latitude, warm-edge populations from extinction.

## INTRODUCTION

The geographic range, encompassing the set of locations where populations of a species occur across the landscape, is a fundamental unit of ecology and biogeography. Understanding how and why abundance varies across the range, and why abundance drops to zero beyond range edges, is relevant for a wide variety of problems, from explaining rarity to forecasting range shifts. Because abundance is the net result of demographic processes such as recruitment, survival, and reproduction, spatial variation in abundance must result from spatial variation in at least some of these vital rates and their combined effects on population growth.

Many hypotheses to explain variation in abundance across species' ranges are predicated on the assumption that a geographic range is a spatial expression of a species' ecological niche, indicating a failure of populations to adapt to conditions at and beyond range edges (1). For example, the classic ‘abundant center hypothesis’ posits that vital rates, population growth, and abundance peak in optimal habitat at the geographic center of the range and decline towards range edges (2). However, empirical support for the abundant center hypothesis is mixed (3–5), likely because spatial and environmental gradients can be decoupled (6, 7), such that environmental optima need not be at the exact geographic center and range-edge populations might occupy patches of optimal habitat. Furthermore, vital rates can respond differently to the same spatial or environmental gradient and need not all decline towards range edges. For example, survival might decrease with temperature while fecundity increases, a phenomenon called demographic compensation (8). If demographic compensation were complete, then population growth would be invariant across the geographic range. Even incomplete compensation could increase the range of environments over which populations can succeed and decrease spatial variation in population growth compared to populations without compensatory changes in vital rates (Fig. 1). Demographic compensation may rescue populations from extinctions due to climate change, at least until reaching a tipping point, beyond which all vital rates decrease and populations crash (9).

**Figure 1.**
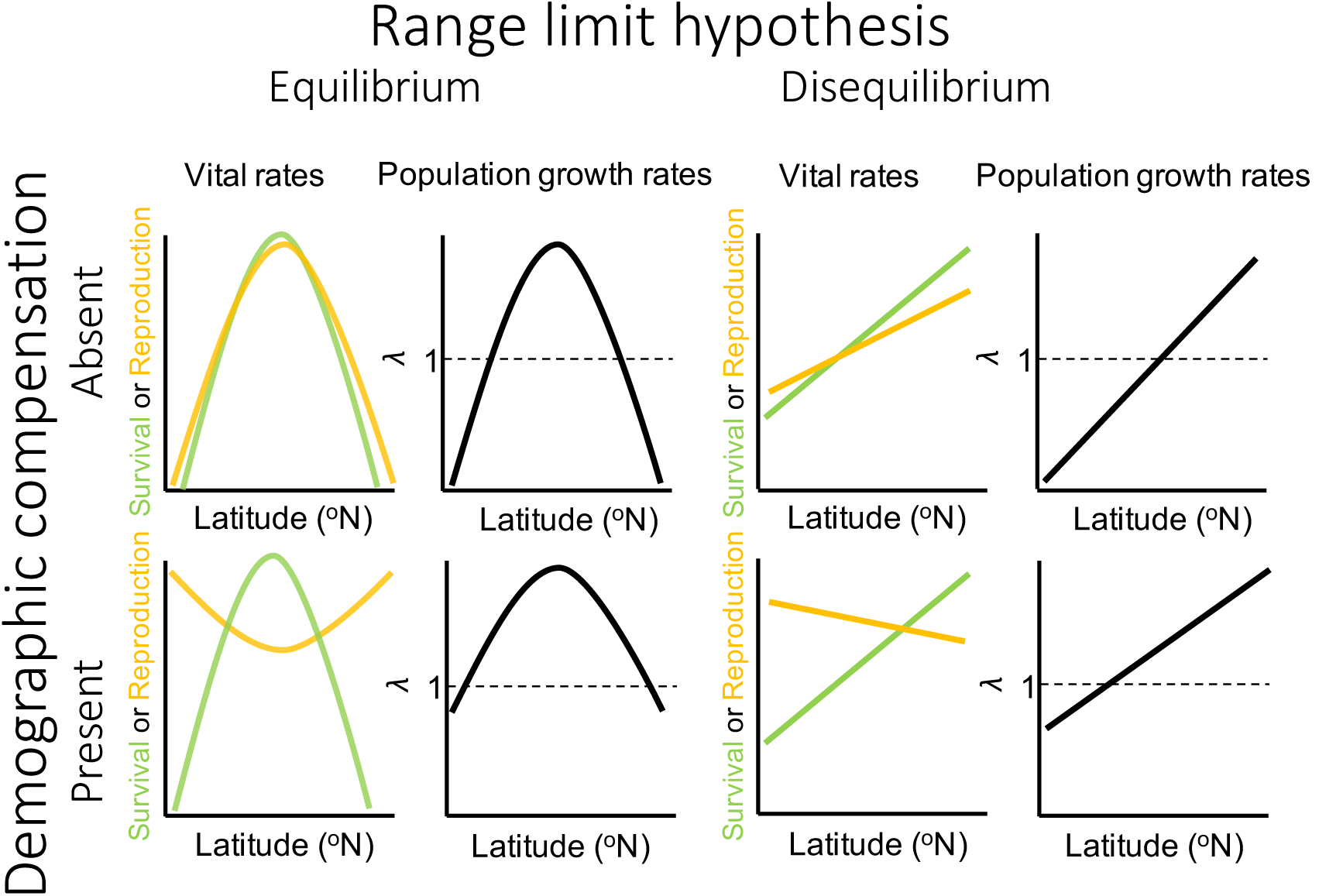
Predictions of how vital rates (the probabilities of survival and reproduction) and population growth rates (*λ*) should vary with latitude in the absence and presence of demographic compensation, under hypotheses of equilibrium and disequilibrium between the species' range and niche. Life history theory predicts negative correlations between vital rates such that fast life history strategies (e.g., high fecundity but low survival or growth from one year to the next) are favored at low latitudes, whereas slow life history strategies (e.g., low fecundity but high survival or growth from one year to the next) are favored at high latitudes.

Whether compensatory or not, spatial variation in vital rates could be driven by phenotypic plasticity or life history evolution. Life history variation can be summarized by a fast-slow continuum along which “fast” life histories have rapid development, high fecundity, reduced longevity, and short generation times while “slow” life histories have delayed development, low fecundity, high longevity, and long generation times (10). Life history theory predicts that selection should favor increased allocation to fast life history characteristics when survival is low or unpredictable, particularly of adults relative to juveniles (11–14). For delayed reproduction to be favored, the potential increase in fecundity by older, larger individuals has to more than offset the risk of mortality until the next opportunity for reproduction. In plants, populations and species from low latitudes are more frequently annual (15) and may reproduce at an earlier age than those from high latitudes (16), creating a fast-slow continuum from low to high latitudes. If demographic compensation is driven by life history evolution, then we might expect to see compensatory negative correlations in vital rates that follow the tradeoffs predicted by the fast-slow continuum (Fig. 1). In particular, we expect that low-latitude populations will exhibit faster life history characteristics than high-latitude populations across a plant species' range (Fig. 1). An outstanding question is whether range and niche limits reflect inadequate compensatory life history evolution along environmental gradients, causing population growth rates to fall below replacement levels at the edges of species' ranges.

Although niche-based mechanisms can define potential abundance and distribution, explaining realized abundance and distribution might require a more dynamic view that accounts for temporal environmental variation and time lags in biological responses to such variation. It is well documented that ranges expand, contract and shift over time, creating ‘leading’ range edges where populations are expanding into newly suitable habitat and ‘trailing’ range edges where populations are contracting from newly unsuitable habitat (17, 18). Disequilibrium between the environment and ranges arises because adaptation, demography and dispersal are usually slower than rates of environmental change over a range of time scales, from glacial-interglacial periods (19) to anthropogenic climate change in recent decades (20). This dynamic view of ranges predicts a linear relationship between vital rates or population growth rate and range position, from low vital rates and declining populations at the trailing edge to high vital rates and stable or growing populations at the leading edge (Fig. 1). It also predicts that demographic compensation will be insufficient to rescue populations at the trailing edge.

In this study, we examine spatial variation in vital rates and projected population growth rates (*λ*) across the geographic range of the scarlet monkeyflower, *Erythranthe cardinalis* (formerly *Mimulus cardinalis). Erythranthe cardinalis* is a perennial herb with a well-described and extensively protected distribution in western North America that spans 12 degrees in latitude across a broad climatic gradient (Fig. 2). Our specific objectives were as follows: 1) Examine how vital rates and *λ* vary range-wide; 2) Determine which vital rates drive variation in *λ*; and 3) Test whether demographic compensation among vital rates buffers spatial variation in *λ*.

**Figure 2.**
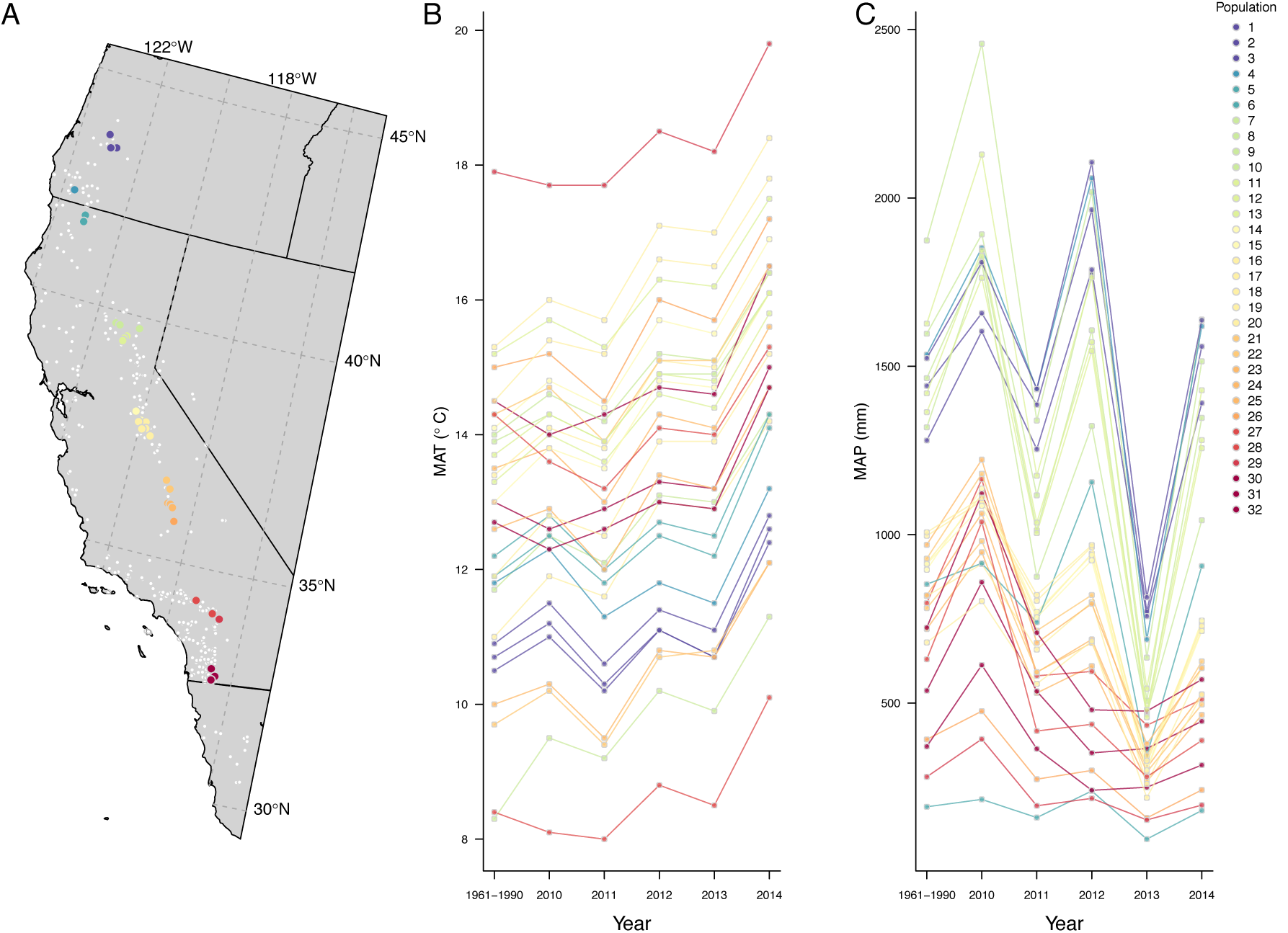
(A) Map of 32 study populations across the geographic distribution of the scarlet monkeyflower. Populations are colored to scale with latitude, with warm colors corresponding to low latitudes and cool colors corresponding to high latitudes; white points correspond to all known occurrences of *E. cardinalis* (Angert et al., in revision). Relationships between (B) mean annual temperature (MAT) and (C) mean annual precipitation (MAP) with latitude across 32 *E. cardinalis* populations. Climate data are derived from ClimateWNA version 5.30 (64).

## RESULTS

### Spatial variation in *λ* and vital rates

Asymptotic projections of population growth rate (*λ*) ranged from 0.1 to 1.9 (Table S1). Lambda varied linearly across the latitudinal gradient, increasing from the southern edge, where *λ*s were uniformly < 1, towards the northern edge, where most populations were stable or increasing (slope = 0.0665, *P* < 0.001, *R^2^* = 0.29; Table S1, Fig. 3). Similar to *λ*, the probability of recruitment and the growth of medium-sized plants increased linearly from south to north (Table S2, Fig. S1). Though not statistically significant, there was a trend of the probability of flowering decreasing from south to north, particularly for small and medium-sized plants (Table S2, Fig. S1). Two other vital rates had non-linear relationships to latitude. The probability of survival peaked at mid latitudes and declined towards northern and southern latitudes, while mean offspring size increased from mid latitudes towards the northern and southern edges (Table S2, Fig. S1). The number of fruits did not vary with latitude (Table S2, Fig. S1).

**Figure 3.**
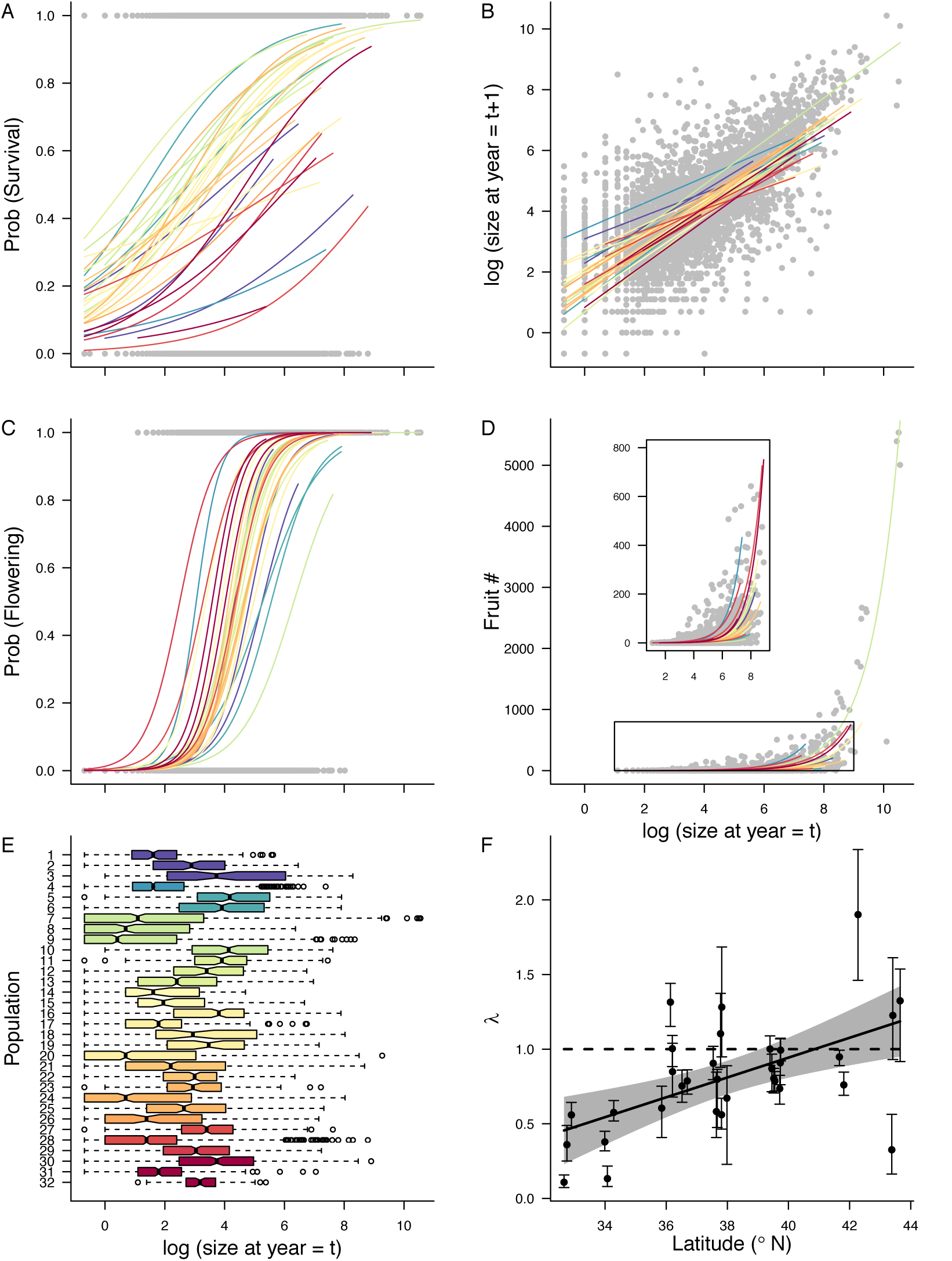
(A-D) Vital rates as a function of size (log-transformed stem length in centimeters). Grey points represent observed values across all populations, and black lines represent model predictions averaged across populations. Colored lines represent model predictions based on population-specific coefficients. Inset panel for (D) shows the same data and fitted lines but omitting individuals with > 800 fruits. (E) Population-specific size distribution of individuals at year = *t.* (F) Population growth rate *λ* as a function of latitude, with vertical bars representing bias-corrected 95% confidence intervals. Dashed line at *λ* = 1 indicates stable population growth, and grey shading corresponds to 95% confidence intervals for predicted *λ* values from linear model.

### Global Life Table Response Experiment

A generalized additive model parameterized by simulated datasets of vital rate coefficients (Appendix S1) accounted for 92.3% of the variation in log (*λ*). Variance in growth and survival probability explained most of the variation in log (*λ*) (44.35% and 34.44%, respectively). Much of the remaining variation in log (*λ*) was explained by variance in recruitment probability (13.74%). Probability of flowering (0.50%), number of fruits (3.00%), number of seeds per fruit (0.55%), and the size distribution of offspring (3.42%) explained the little remaining variation.

### Demographic compensation

Of 21 possible pairwise correlations among population-specific vital rate contributions, 4 were significantly negative (*P* < 0.05; Table 1, Fig. S2). Specifically, there were negative correlations between contributions of survival and flowering probabilities, survival probability and number of fruits, number of fruits and recruitment probability, and number of fruits and the size distribution of offspring (*P* < 0.05; Table 1, Fig. S2). The observed proportion of negative correlations was significantly greater than expected by chance (P = 0.0186), consistent with significant demographic compensation.

**Table 1.** Pairwise Spearman rank correlations among vital rate contributions.

|  | Prob<br>(Survival) | Growth | Prob<br>(Flowering) | Fruit number | Seeds<br>per fruit | Prob<br>(Recruitment) | Distribution of<br>offspring size |
| --- | --- | --- | --- | --- | --- | --- | --- |
| Prob (Survival) |  | 0.17 | -0.42<sup>**</sup> | -0.47<sup>**</sup> | 0.05 | 0.37 | -0.18 |
| Growth |  |  | -0.22 | 0.17 | -0.07 | 0.20 | -0.24 |
| Prob (Flowering) |  |  |  | 0.61 | 0.36 | -0.21 | -0.13 |
| Fruit number |  |  |  |  | 0.11 | -0.38<sup>*</sup> | -0.30<sup>*</sup> |
| Seeds per fruit |  |  |  |  |  | -0.26 | -0.23 |
| Prob (Recruitment) |  |  |  |  |  |  | -0.12 |

\* $P < 0.05$ ; \*\* $P < 0.01$

Population-specific vital rate contributions to *λ* varied with latitude (Table S3, Fig. 4). Contributions of survival probability and fruit number were unimodal with respect to latitude. Survival contributions peaked at mid latitudes, where survival rates increased *λ*, and decreased towards the north and south, where survival rates decreased *λ* (Table S3, Fig. 4). Fruit number showed a weaker but opposite pattern with the largest negative contributions at mid latitudes and positive contributions in the north and south (Tables S3, Fig. 4). The contributions of growth and recruitment probability increased from negative values in the south to positive values in the north, while the contribution of flowering probability decreased from positive values in the south to negative values in the north (Table S3, Fig. 4). Contributions of offspring size did not vary with latitude (Table S3, Fig. 4). Consistent with the global life table response experiment based on GAM, the overall magnitude of population-specific vital rate contributions to *λ* was highest for survival probability and growth (Fig. 4a, b), and lowest for flowering probability (Fig. 4c), indicating that the positive contributions of flowering probability were too low to bolster *λ* in southern populations.

**Figure 4.**
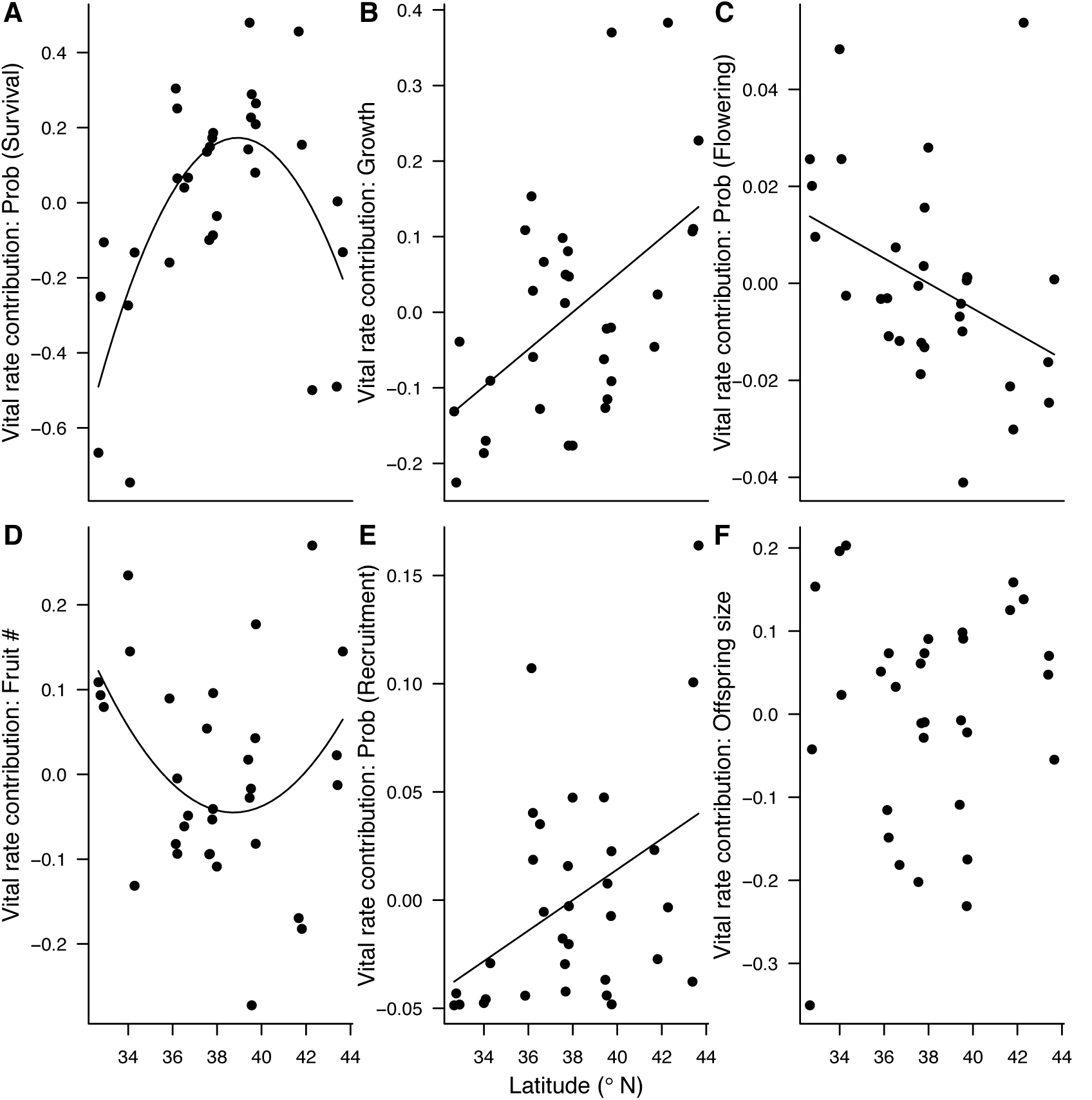
Latitude vs. population-specific contribution of each vital rate. Solid lines represent slopes that are greater than 0 at *P* < 0.05.

## DISCUSSION

Demographic compensation has been proposed as a mechanism for buffering warm-edge populations from climate change, at least until reaching a tipping point (9), yet few studies have examined the role of demographic compensation and life history shifts in preventing population collapse. Consistent with disequilibrium between the range and niche, we found that population growth rates increased with latitude, suggesting that low-latitude trailing edge populations are contracting from newly unfavorable environments, whereas high-latitude leading edge populations are increasing in recently ameliorated areas. Differences in growth, survival, and recruitment drove spatial variation in *λ*, with growth and recruitment probabilities increasing with latitude and survival probabilities decreasing from mid latitudes towards the northern and southern edges. The presence of statistically significant demographic compensation among vital rates indicates that vital rates may respond individualistically to the same environmental gradient and that single vital rates may fail to predict population performance. However, although there was demographic compensation, small positive contributions from a greater probability of flowering and greater fruit number did not buffer southern populations from the large, negative effects of low survival, growth, and recruitment. Below, we place our key findings in the context of life history strategies and past and ongoing climate change.

### Disequilibrium between range and niche limits

Our results contrast with a large body of work suggesting that range limits often coincide with niche limits. Reciprocal transplant experiments of several species reveal a decrease in performance beyond range edges, and provide weak support for reduced performance at range edges (21–23). Supporting the hypothesis that fitness limits species' ranges along elevation gradients, a transplant experiment of *E. cardinalis* within and beyond its elevation range revealed severely reduced fitness at and beyond the upper elevation edge (24). Further, experimental and observational studies of other plant species have shown reductions in abundance and fecundity at or beyond range limits (25–27), and declines in population growth rates at or beyond range margins (28, 29). As ongoing climate change continues to diminish equilibrium between ranges and niches, recent studies such as the present one may reveal stronger demographic signatures of disequilibrium than prior work.

Reflecting such disequilibrium, populations at the southern edge behaved differently than populations at the northern edge. Most northern edge populations were either stable or growing, perhaps due to an influx of pre-adapted genotypes from more southerly populations or amelioration of growth constraints in northern populations. The net result to date suggests a “lean” range shift (30, 31), where range limits have remained stable but the central tendency of the distribution is moving northward within the existing range. A lean range shift potentially reflects disequilibrium with climate at both the leading and trailing edges, with disequilibrium at leading edges involving lags in colonization and adaptation, and disequilibrium at trailing edges resulting from delays in extinction (32). Southern edge populations were indeed declining, perhaps in areas that were historically suitable. California experienced severe, record-setting drought conditions from 2012 through 2014 (33, 34), compounded by record high temperatures (35; Fig. 2b), in 3 of the 4 yearly transitions in the present study. High-latitude populations of *E. cardinalis* have already begun to experience temperatures found historically at the latitudinal range center (Fig. 2b). Interestingly, one northern population with extremely high estimated population growth (population 4, Table S1) is in unusually unstable substrate and exhibits annualized life history characteristics that are more typical of southern populations, perhaps providing a window into the potential population dynamics of southern populations under more favorable conditions. In the present study, high mortality in southern populations in recent drought years reduced sample sizes for vital rate models to parameterize yearly integral projection models, requiring pooling of data across multiple yearly transitions. As we continue to monitor these populations over time, we will gain a better understanding of the impacts of recent climate change on population dynamics by linking yearly variation in weather to yearly variation in vital rates and population growth rates. Further supporting disequilibrium between the range and niche, previous studies of demography of *E. cardinalis* at mid latitudes found higher *λ*s at the upper elevation edge when compared to mid elevations (36, 37), and a recent transplant study documented that As of populations transplanted beyond the northern edge were greater than 1, comparable to those transplanted within the range (38). Consistent with our finding of high mean fitness in northern edge populations, previous transplant studies of a coastal dune plant with a similar latitudinal range as *E. cardinalis* reveal a pattern of population mean fitness increasing towards and beyond the northern range edge (39, 40). However, population growth rates do not always vary predictably with latitude (7, 9), likely due to local environmental factors and/or demographic compensation.

### Inadequate demographic compensation

Demographic compensation may buffer populations at low latitudes against extinction risks associated with climate change (9). In this study, increases in the probability of flowering and fruit number in low-latitude populations (indicated by small, positive contributions to variation in *λ*) suggest that population growth rates would likely be even lower at the southern edge in the absence of demographic compensation. However, even in the face of significant demographic compensation across the geographic range of *E. cardinalis*, population mean fitness was substantially reduced in low-latitude populations relative to mid- and high-latitude populations. The magnitude of compensatory increases in reproductive vital rates in low-latitude populations was too small to offset the large, negative contributions of survival, growth, and recruitment (Figs. 4, S1, S2). In contrast, in high-latitude populations, low probabilities of survival and flowering were offset by high growth and recruitment (Figs. 4, S1, S2), thus promoting population growth.

### Latitudinal gradient in life history strategy

Spatial variation in vital rates and population-specific vital rate contributions are consistent with life history theory and with life history variation observed in a common garden study of *E. cardinalis* populations sampled across its latitudinal range (41). Matching expectations from life history theory (12, 15, 42, 43), low-latitude populations of *E. cardinalis* experience greater inter-annual variation in precipitation (41), germinate, photosynthesize and grow faster in a common garden (41), and exhibit a faster, more annualized life history strategy in nature (this study), whereas high-latitude populations from more temporally stable environments grow more slowly (41) and are uniformly perennial (this study). In particular, plants at low latitudes tend to grow quickly within a growing season, flower once, and then die (supported by low growth and survival but high flowering probabilities; Fig. S1), whereas plants at high latitudes grow slowly within a growing season, do not flower every year, and accumulate more growth from one year to the next (supported by high growth and low flowering probabilities; Fig. S1). Similar geographic and climatic clines in life history traits have been observed in the common monkeyflower (44), common mullein (15), Queen Anne's Lace (16), and purple loosestrife (45). The compensatory demographic variation observed across populations of *E. cardinalis* supports life history theory predicting tradeoffs along the “fast-slow continuum” between slow development, less frequent but multiple bouts of reproduction, and long lifespan versus fast development, high reproduction, and short longevity (46–48). Linking demographic compensation to life history theory allows a broader understanding of how spatial variation in vital rates shapes species' geographic ranges.

### Conclusions

This study highlights the importance of non-equilibrium processes in shaping species' geographic distributions. In contrast with previous work showing that geographic range limits often coincide with niche limits (49), leading-edge populations appear to be expanding in response to contemporary anthropogenic climate change (50, 51). Consistent with a recent poleward shift of the species' climatic niche, *E. cardinalis* populations at the northern range edge have begun to encounter temperatures similar to those occurring historically at central latitudes (Fig. 2b). As species continue to shift their climatic envelopes in response to recent climate change, support for hypotheses assuming equilibrium between range and niche limits may continue to weaken. Climatic tipping points where demographic compensation breaks down, may result in population collapse (9). We show that even statistically significant demographic compensation may not be sufficient to buffer warm, trailing edge populations against extinction. Because individual vital rates responded differently to the same latitudinal gradient, we warn against the use of one or a few vital rates to predict population performance. As additional demographic data accumulate for multiple generations, populations, and species across broad spatial scales and environmental gradients we will gain a more comprehensive understanding of how the environment and geography shape vital rates and in turn population dynamics, allowing for better forecasts of range shifts.

## MATERIALS AND METHODS

### Study system

*Erythranthe cardinalis* (Phrymaceae) is a perennial forb that grows along seeps, streamsides, and riverbanks in western North America. Individuals can spread via rhizomes but recruitment occurs almost exclusively from seeds (Angert, pers. obs.). The species' latitudinal range extends from central Oregon, USA to northern Baja California, Mexico (Fig. 2a). Within this latitudinal extent, populations occur across a broad range of elevations and climates (Table S4; Fig. 2b, c), from sea level up to ca. 2400 m (52). However, latitude and elevation of occurrences covary (Pearson r = -0.57, P < 0.05), such that northern populations are primarily at low elevations while southern populations can extend to higher elevations.

### Demographic surveys

We established long-term census plots in 32 populations spanning almost the full latitudinal extent of the species' range (Fig. 2a, Appendix S1). During August and September 2010, census transects encompassing multiple areas suitable for all life history transitions were established within each population except populations 4, 23, and 26 (Fig. 2a, Table S4), which were added in 2011, 2012, and 2012, respectively. Due to unavoidable variation in microhabitat and plant density, the number and length of transects varied across populations (Table S4). Our aim was to survey at least three transects per population that together encompassed at least 200 individuals and contained habitat suitable for future recruitment. Transects were anchored with rebar, removable bolts drilled by hand into rock slabs, or nails in crevices or tree trunks. New or replacement transects were added as needed to maintain target sample size or when older transects were lost due to flooding or treefall.

Every *E. cardinalis* individual was uniquely identified by (x, y)-coordinates, using the transects as y-axes and perpendicular distances from the transects as x-axes. Some individuals also received uniquely numbered tags to confirm the alignment of the transect from year to year. Not every individual could receive a tag due to microhabitat constraints (e.g. rock slabs) and requirements imposed by permitting agencies. Censuses were conducted each autumn, after most annual reproduction was complete, from 2010 to 2014 to record annual survival, growth, reproduction, and recruitment. In total, the fates of 11, 244 plants were recorded (Table S4).

To estimate size and annual growth for each plant, up to 5 non-flowering and 5 flowering stems were measured from the ground to the base of the last pair of leaves; all remaining stems were tallied and used to estimate total stem length based on the average stem length of the 10 measured stems. Plant reproduction was estimated as the product of the number of mature fruits on up to 5 stems of a given individual times the total number of flowering stems on that individual times the population mean seed number per fruit in a given year. Each fruit may contain 500-2500 tiny seeds that cannot be counted in the field. To estimate seed number per fruit, two fruits were harvested each fall 2010-2012 from up to 10 individuals growing downstream of the census transects, or, when downstream sampling was not possible, several hundred meters upstream of the census transects. For 2010-2011 samples, subsamples of approximately 200 seeds per fruit were counted under a dissecting microscope and weighed to determine the relationship between seed mass and seed number. Seed number per fruit was then estimated from total seed mass, which has been shown to accurately capture true seed number (36). For 2012 samples, seeds were photographed and then counted using image analysis software (ImageJ). Fruits could not be obtained for some population-by-year combinations (we did not obtain any seed counts for 2013), so average seed number per fruit across all other years for that population was used instead. For 8 populations from which we were unable to obtain fruits, we used estimates of seed number per fruit from the geographically closest population from which fruits were collected. Based on experiments to estimate seedling recruitment and seed dormancy (Appendix S1), seed dormancy was set to zero for these analyses and seedling recruitment was estimated by dividing the number of new recruits by total seed production in the prior year.

### Integral projection models

To estimate population growth rates for each study population, we used integral projection models (IPMs), which are analogous to population matrix models but model vital rates as a function of individual plant size, which varies with latitude (Fig. 3a-e) rather than relying on discrete stages (53, 54). First, we pooled data across all populations and years to construct a global model of each of four vital rates (probability of survival, growth, probability of flowering, and fruit number; Table S5) as a function of size (fixed effect), year (random effect), and population (random effect) in the lme4 package (version 1.1-12, 57) in R 3.3.1 (56). We included log-transformed total stem length in year *t* as the size measurement in all vital rate models, and compared models with and without size as a predictor variable based on Akaike information criterion for small samples (AIC_c_). Size was a significant predictor of all four vital rates. For each vital rate, we compared full models with random slopes and intercepts for both population and year to progressively simpler models with the simplest model including random intercepts for population and year. We retained those terms that were significant in log-likelihood ratio tests (*P* < 0.05; Table S5). We extracted population-specific coefficients for each vital rate function to parameterize the IPM for each population (Table S1). Due to small sample sizes in some populations in some years, we constructed the IPM kernel for a given population using data pooled across all years (Appendix S1).

We discretized IPM kernels into a matrix with 100 size bins, with size ranging from 0.9 times the minimum and 1.1 times the maximum size observed in year *t* or year *t* + 1 in each population. To correct for the “eviction” of individuals falling beyond this size range (57), we assigned individuals to the smallest size bin in the case of offspring, and to the largest size bin in the case of large adults (58). We quantified population growth rate (*λ*) as the dominant eigenvalue of the matrix. We performed all analyses in R v. 3.3.1 using code modified from appendices in Merow et al. (58) and Rees et al. (59). To obtain 95% confidence intervals around *λ* estimates for each population, we bootstrapped the data 2000 times, allowing for assessments of whether the *λ* estimate for each population was statistically greater than, less than, or not different from 1 (Appendix S1).

### Analysis of latitude vs. *λ* and vital rates

To assess how *λ* varies across latitude, we used linear regressions with *λ* as the response variable and latitude as the predictor variable. We compared models with and without quadratic terms and used AIC to select the best fitting models. For the vital rates that varied with size (survival, growth, probability of flowering, and number of fruits), we first divided individuals in each population among small (0-20% quantile), medium (40-60% quantile), and large (80-100% quantile) size classes, which varied depending on each population's overall distribution of individual sizes. We then estimated population- and size-specific probability of survival, growth, probability of flowering, and number of fruits. Survival probability is the proportion of individuals that survived in each size class at each population, growth is the mean size in year *t* + 1 across all individuals of each size class in each population, flowering probability is the proportion of individuals that flowered in each size class in each population, and number of fruits is the mean number of fruits in year *t* + 1 across all individuals of each size class in each population (small size class was omitted from models of fruit number because “small” plants only flowered at 3 populations). We regressed mean vital rates for each size class against latitude, with and without a quadratic term for latitude, and used AIC to assess whether to include the quadratic term or not. Because the study populations spanned ~1700 m in elevation, we initially performed similar analyses with elevation as a predictor variable but found no statistically significant relationships between elevation and vital rates or lambda, and models including latitude alone performed better than any models including latitude and elevation. Thus, here we only present results including latitude as a predictor variable.

### Global Life Table Response Experiment

To identify which vital rates most contribute to observed differences in *λ* among populations, we initially performed a standard life table response experiment (60), but the range of variation in parameter values among populations resulted in a poor linear approximation of *λ* as a nonlinear function of the parameters that vary among populations. Instead, we fit a generalized additive model (GAM) using the ‘gam’ function in the mgcv package (version 1.8-12;, 61), with log (*λ*) as the response variable and smoothed functions of vital rate parameters as explanatory variables (62, 63 Appendix S1). To obtain contributions at the level of each vital rate as a whole (rather than each parameter in each vital rate function), we summed across all coefficients of a given vital rate (e.g., the survival contribution equals the sum of survival slope contribution and the survival intercept contribution). Vital rate contributions to variability in log (*λ*) were normalized to sum to 100%.

### Demographic compensation

Following Villellas et al. (8), we examined demographic compensation by testing for negative correlations among vital rates, weighted by their population-specific contributions to variation in *λ* (Appendix S1). To test whether there are more negative correlations among vital rate contributions than expected by chance, we obtained Spearman rank correlations between all pairs of vital rate contributions. We then determined the observed percentage of correlations that were significantly negative (*P* < 0.05) based on a one-tailed test. Next, in each of 10,000 iterations, we randomly permuted contributions for each vital rate among populations, calculated Spearman rank correlations, and determined the percentage of significantly negative correlations. Thus, we obtained a null distribution of percentages of negative correlations against which to compare our observed percentage. We inferred statistically significant demographic compensation based on the proportion of values in the null distribution that were greater than or equal to the observed percentage of negative correlations (8). In theory one could also test whether there are fewer positive correlations than expected by chance, but there were no significantly positive correlations among vital rate contributions in the observed data. To assist in our interpretation of the test of demographic compensation, we also examined how vital rate contributions varied with latitude. For each vital rate contribution, we used linear regressions with and without quadratic terms, and then used AIC to select the best fitting models with each vital rate contribution as the response variable and latitude as the predictor variable.

## ACKNOWLEDGEMENTS

ALA conceived of the study; ALA led data collection and curation; SNS analyzed the data; SNS and ALA wrote the manuscript. We thank A. Agneray, M. Bayly, P. Beattie, M. Bontrager, B. Econopouly, C. Fallon, B. Gass, K. Hafeez, E. Hinman, Q. Li, J. Perce, D. Picklum, A. Rosvall, J. Smith, L. Super, M. Wiebush, and A. Wilkinson for contributing to data collection in the lab and/or field. Barb Gass and C. Van Den Elzen assisted with data management. We thank J. Williams for discussions and feedback about IPMs, and S. Ellner for advice and generously sharing code to simulate parameter datasets for the GAM. The Angert lab (especially M. Bayly, M. Bontrager, R. Germain, and J. R. Paul) and the Ackerly lab (especially M. Oldfather) provided feedback on earlier versions of this work. This work was funded by National Science Foundation DEB-0950171 and 1112837 to ALA and J. R. Paul, and SNS was supported by NSF DEB-1210879 and DBI-1523866.

## SUPPORTING INFORMATION

### Appendix S1 Supplementary methods

#### Selection of study populations

From north to south, we surveyed populations from the following regions: (1) central Oregon, (2) southern Oregon - northern California, (3) northern Sierra Nevada Mountains, (4) central Sierra Nevada Mountains, (5) southern Sierra Nevada Mountains, (6) Transverse Ranges, and (7) southern California (Table S4, Fig. 2a). Region 1 includes the northernmost extant populations that we located after repeated field surveys from 2007-2010. Region 7 does not encompass the southernmost populations that have been recorded because logistical constraints prevented us from working in Baja California, Mexico, where there are disjunct occurrences spanning ~160 km from north to south (<10% of the overall latitudinal gradient; Fig. 2a). However, study populations in southern California (region 7) occur in similar habitat as reported on herbarium specimens from Baja California (e.g., seeps and streamsides), and the inclusion of herbarium specimens from Baja California increases the range of mean annual temperature and mean annual precipitation among recorded populations by less than 0.1% relative to the range of climates when excluding specimens from Baja California based on climate variables for 1961-1990 and 2013 derived from ClimateWNA version 4.62 (64). We oriented the study transect through the mountainous Sierra Nevada regions because there are extensive tracts of protected public lands and so that we could nest replicate elevation transects within several latitudinal regions. Accordingly, within each Sierra Nevada region (regions 3-5), we selected 6 populations at three elevations per each of two replicate drainages. Outside of the Sierra Nevada regions (regions 1-2, 6-7), we selected 3 populations in multiple drainages (Table S4, Fig. 2a). Populations were selected based on habitat quality and accessibility during extensive field surveys guided in part by collection records from regional herbaria.

#### Seed transitions

In a subset of 7 populations, we created seed addition plots for estimating seedling recruitment and seed dormancy. Within each population, at each of 10 20-cm × 20-cm plots placed haphazardly within suitable microhabitat for germination, we added 100 seeds in September 2011. Each seed addition plot was paired with a no-seed-addition control. To prevent seed entry and escape, we covered the plots in fine mesh (“No Thrips”, 150 × 150 μ opening size, Green-Tek, Inc., Edgerton, WI, USA) each fall and winter and removed it after floodwaters receded every spring. Plots were censused each spring and fall for 3 years. Although a prior study at mid latitudes demonstrated that a small fraction of *E. cardinalis* seeds can survive in the seed bank and germinate in their second year (36), during this study period we only observed germination in the first spring after seed addition and none at any subsequent census. Thus, seed dormancy was set to zero for these analyses and seedling recruitment was estimated by dividing the number of new recruits by total seed production in the prior year.

##### Integral projection models

For each population, we created a kernel *K*(*Z*', *Z*) describing how size *z* of individuals in year *t* determines size *z'* of individuals in year *t*+1 (54, 58):

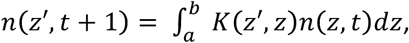

where *n*(*z*, *t*)*dz* represents the number of individuals whose size falls within the range [*z*, *z* + *dz*]. We divided this IPM kernel into survival/growth *(P)* and fecundity *(F)* kernels:

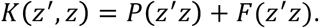

The survival/growth kernel is:

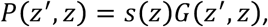

where *S*(*Z*) is the size-dependent probability of survival and *G*(*Z*',*Z*) is size-dependent growth or shrinkage. The fecundity kernel is:

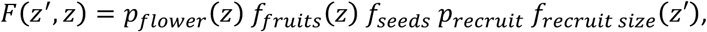

where *P_flower_*(*z*) is probability of flowering as a function of size in year *t, f_fruits_(.z)* is fruit number as a function of size in year t, *f_seeds_* is the average number of seeds per fruit, *p_recruit_* is the probability of recruitment, and *f_recruit size_*(*.z*) is the size distribution of new recruits in year *t*+ 1.

##### Bootstrapping to obtain confidence intervals for λ

We sampled unique individuals from each population with replacement to create replicate bootstrap datasets, where the size of each population's bootstrap was equal to the number of unique individuals in original dataset. Subsequently, we ran vital rate models as described above on each bootstrapped dataset to obtain population-specific coefficients to parameterize population-specific IPM kernels. We then ran IPMs as described above on each bootstrapped dataset for each population and re-calculated, allowing for assessments of whether the *λ* estimate for each population was statistically greater than, less than, or not different from 1.

##### Global Life Table Response Experiment

Because of low sample size (response variable of *λ* for *N* = 32 populations but 12 explanatory variables corresponding to vital rate parameters that vary among populations), we first simulated 10,000 sets of vital rate parameters (62). To do so, we generated a large set of parameters with the same pairwise Spearman rank-correlations and marginal distributions as the observed parameters (65). We then used our existing IPM framework to estimate *λ* for each of these 10,000 simulated parameter sets, using the full size range observed across all populations rather than population-specific size ranges. The resulting and parameter values were used in GAM. We evaluated the accuracy (R^2^) of the GAM at our observed parameter values (S. Ellner, *pers. comm.*). We decomposed the variance in *λ* by extracting fitted values of each model term from GAM fit to observed parameter estimates, and then obtaining the weighted variance-covariance matrix of those terms (63). We assessed the proportion of variance in *A* explained by variance in each parameter acting on its own by dividing each diagonal value of the weighted variance-covariance matrix by the sum across all diagonal values.

##### Demographic compensation

First, we calculated *λ* for a reference IPM based on parameter estimates averaged across all 32 populations. Next, we perturbed each of 13 vital rate coefficients in turn by adding a small value (0.01). We re-calculated *λ* for the reference IPM after each perturbation, and estimated sensitivity for each coefficient as the difference between perturbed *λ* and unperturbed *λ*, divided by the amount of perturbation (Table S6). Subsequently, we determined the contribution *C_i,n_* for vital rate parameter *i* and population *n* as *C_i,n_ =* (*v_i,n_ – v_i,r_*)×*S_i,r_*, where *v_i,n_* and *v_i,r_* correspond to vital rate *i* for population *n* and the reference population, respectively, and *S_i_*_,*r*_ is the sensitivity of *λ* calculated from the reference matrix to perturbations in vital rate *i* (8). To obtain contributions for each vital rate as a whole (rather than each vital rate parameter), we summed across all coefficients of a given vital rate as described above.

#### Appendix S2 Supplementary tables

**Table S1.**
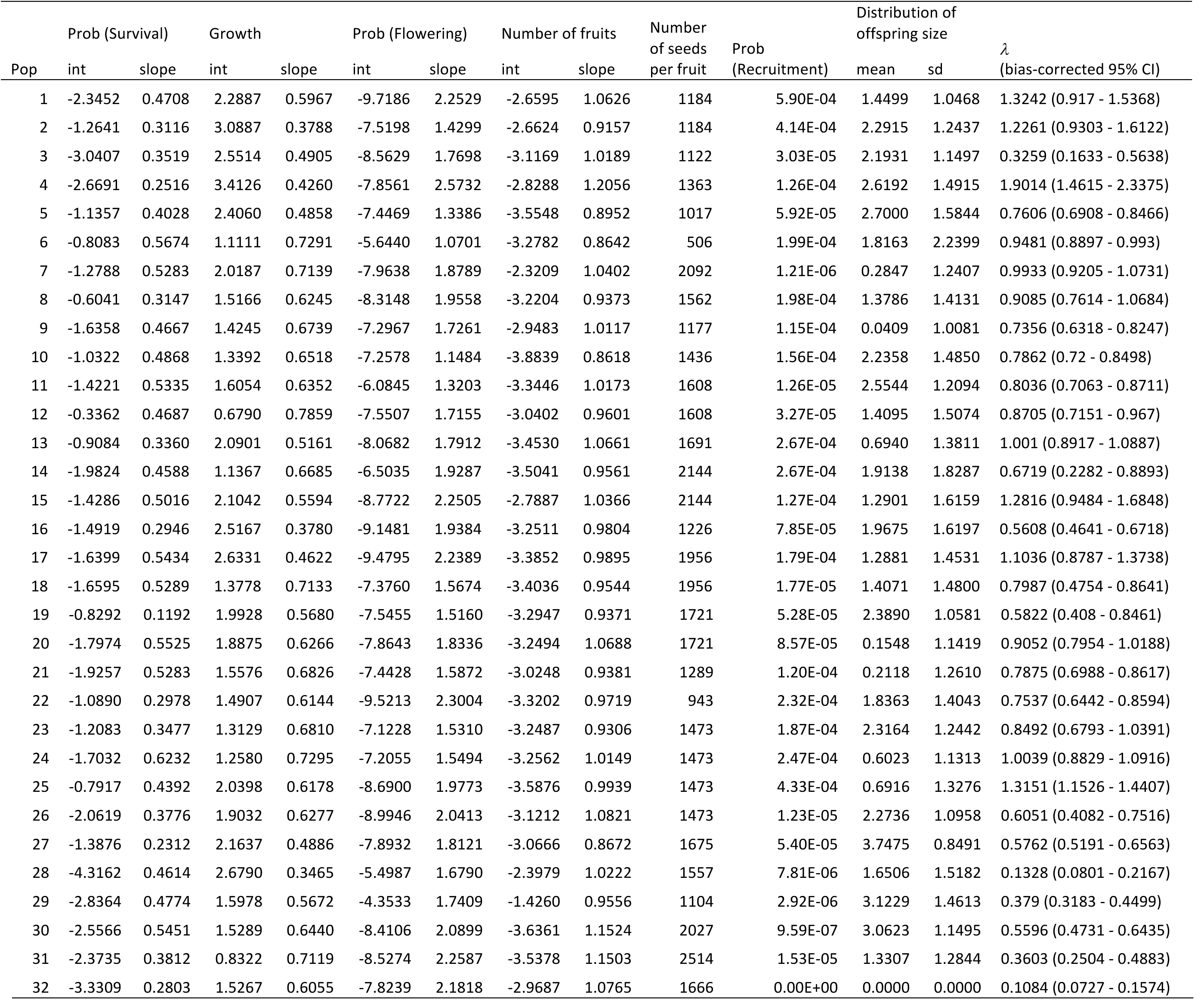
Vital rate coefficients used to parameterize IPM for each *E. cardinalis* study population (population codes are described in Table S1). A constant standard deviation for growth, 1.1014, was used across all populations.

**Table S2.** Regression coefficients and adjusted R^2^ for models of vital rates vs. latitude. Mean fruit number was log-transformed and only large plants were included because small & medium plants did not make many fruits. Growth (measured as size in year *t*+1) and offspring size were also log-transformed. Recruitment probability and offspring size are population-level measurements and do not vary among individuals within a population. In models with quadratic terms marked with “-“ instead of a coefficient estimate, quadratic terms were dropped based on model selection using AIC.

|  |  | Prob (Survival) | Growth | Prob (Flowering) | Fruit number | Prob<br>(Recruitment) | Offspring size |
| --- | --- | --- | --- | --- | --- | --- | --- |
| small | Latitude | 4.29E-01<sup>+</sup> | -3.21<sup>**</sup> | -1.88E-03<sup>+</sup> | - | - |  |
|  | Latitude<sup>2</sup> | -5.48E-03<sup>+</sup> | 4.31E-02<sup>**</sup> |  | - | - |  |
| | Adj. $R^2$ | 0.07 | 0.24 | 0.06 | - | - | |
| medium | Latitude | 5.32E-01<sup>*</sup> | 1.41E-01<sup>**</sup> | -3.78E-01<sup>+</sup> | - | - |  |
|  | Latitude<sup>2</sup> | -6.72E-03<sup>+</sup> | - | 4.76E-03<sup>+</sup> | - | - |  |
| | Adj. $R^2$ | 0.14 | 0.20 | 0.11 | - | - | |
| large | Latitude | 1.10<sup>***</sup> | 2.23E+00<sup>+</sup> | -1.71E-02 | -1.38E-02 | 1.96E-05<sup>*</sup> | -2.93<sup>*</sup> |
|  | Latitude<sup>2</sup> | -1.42E-02<sup>***</sup> | -2.79E-02<sup>+</sup> |  | - | - | 0.04<sup>*</sup> |
| | Adj. $R^2$ | 0.32 | 0.16 | 0.05 | -0.03 | 0.15 | 0.12 |

<sup>+</sup>P < 0.1; <sup>\*</sup>P < 0.05; <sup>\*\*</sup>P < 0.01; <sup>\*\*\*</sup>P < 0.001

**Table S3.** Regression coefficients and adjusted R^2^ for models of population-specific vital rate contributions (following VilleMas et al. 2015) vs. latitude. Quadratic terms marked with “-” instead of a coefficient estimate represent models in which quadratic terms were dropped based on model selection using AIC.

|  | Prob<br>(Survival) | Growth | Prob<br>(Flowering) | Fruit # | Prob<br>(Recruitment) | Offspring<br>size |
| --- | --- | --- | --- | --- | --- | --- |
| Latitude | 1.31<sup>***</sup> | 2.46E-02<sup>**</sup> | -2.59E-03<sup>*</sup> | -3.51E-01<sup>*</sup> | 7.06E-03<sup>*</sup> | 5.15E-03 |
| Latitude<sup>2</sup> | -1.69E-02<sup>***</sup> | - | - | 4.53E-03<sup>*</sup> | - | - |
| Adj. $R^2$ | 0.43 | 0.24 | 0.12 | 0.12 | 0.15 | -0.02 |

<sup>\*</sup>P < 0.1; <sup>\*</sup>P < 0.05; <sup>\*\*</sup>P < 0.01; <sup>\*\*\*</sup>P < 0.001

**Table S4.**
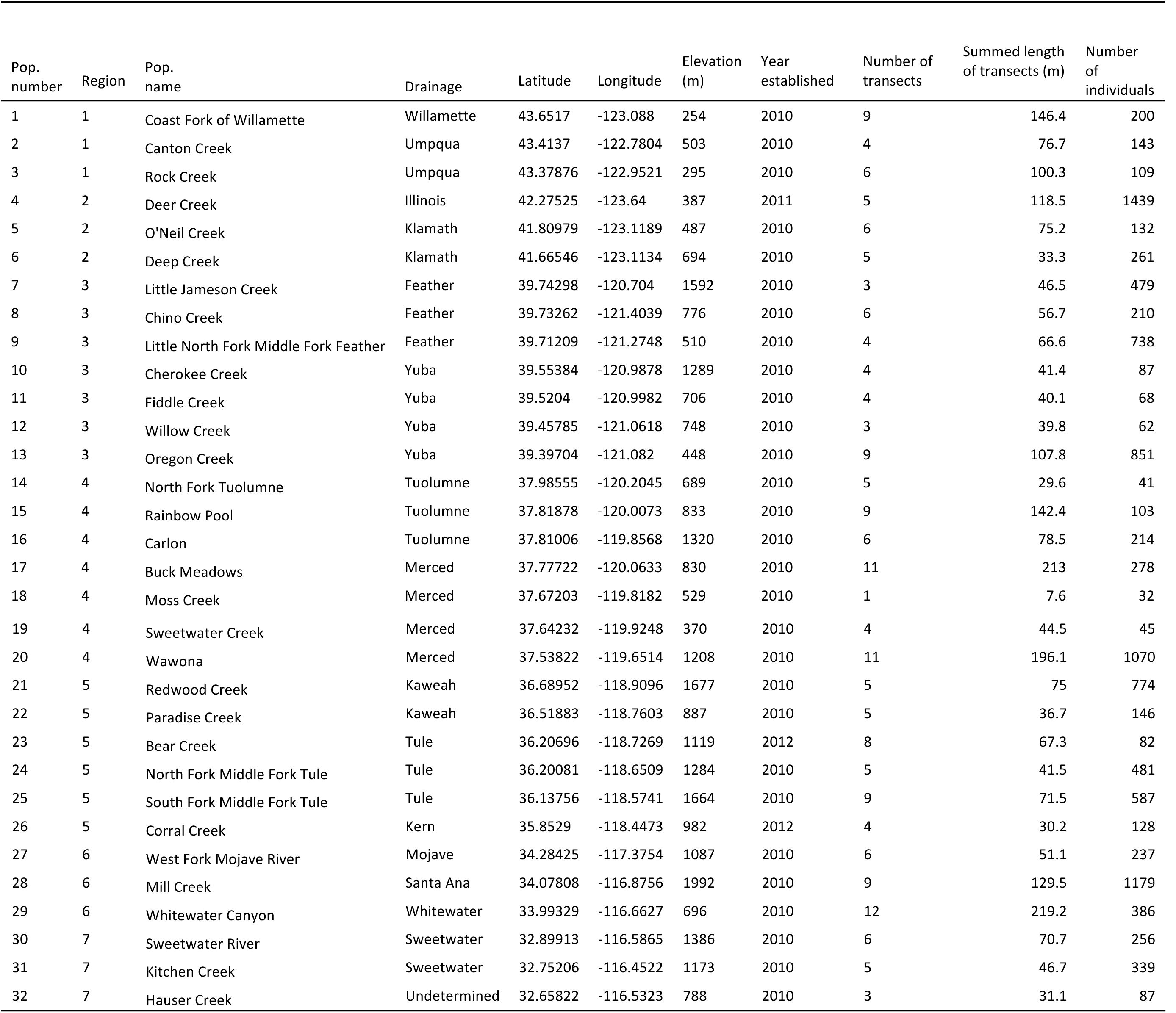
Geographic and sampling information for *E. cardinalis* populations. The number and length of transects for each population are cumulative over time, rather than per year, because plots were abandoned for various reasons or added in subsequent years. Thus, in any given year the sampled area could be less than the cumulative number and length of transects reported here.

**Table S5.**
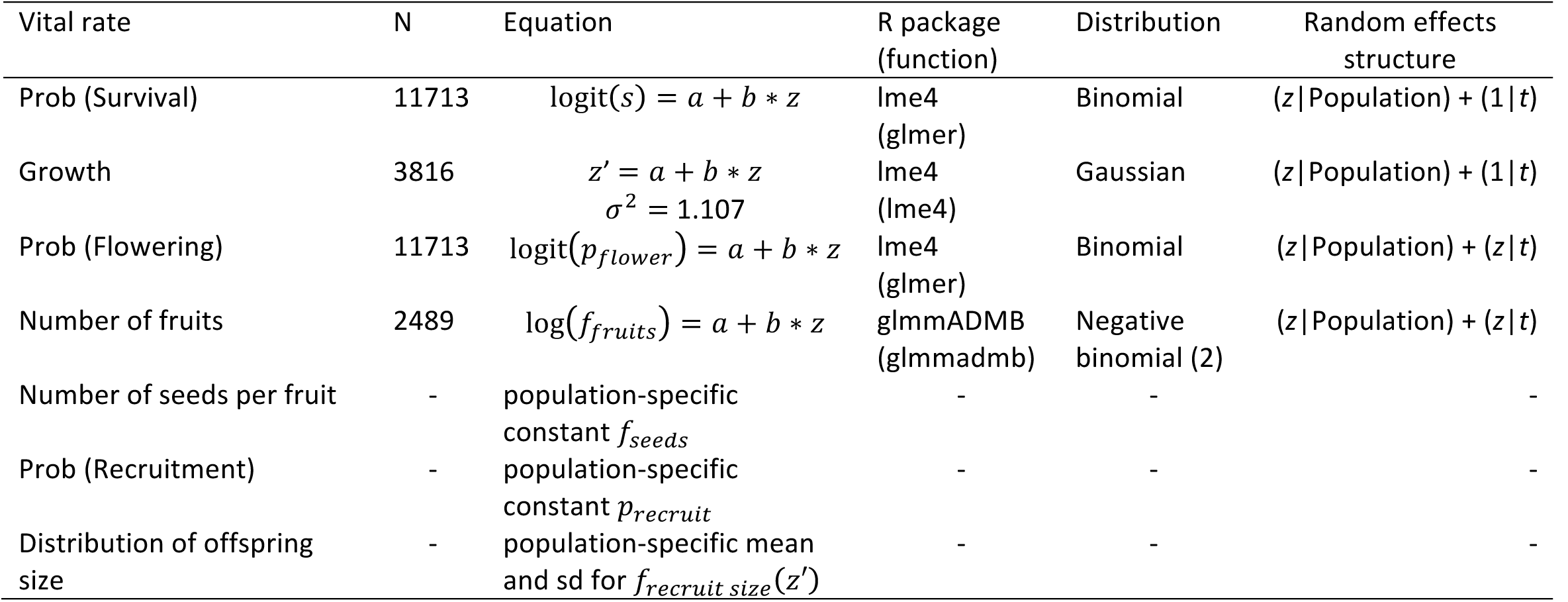
Vital rate functions used in integral projection models and global life table response experiment.

**Table S6.** Sensitivities and *λ* estimates derived from reference matrix used to estimate population-specific vital rate contributions. Unperturbed *λ* from reference matrix = 0.8527.

| Vital rate | Parameter | Sensitivity | Perturbed $\lambda$ |
| --- | --- | --- | --- |
| P(Survival) | intercept | 0.3056 | 0.8558 |
| P(Survival) | slope | 1.2251 | 0.8649 |
| Growth | intercept | 0.4978 | 0.8577 |
| Growth | slope | 2.3650 | 0.8763 |
| Growth | SD | 0.3996 | 0.8567 |
| P(Flowering) | intercept | 0.0158 | 0.8529 |
| P(Flowering) | slope | 0.0732 | 0.8534 |
| Number of fruits | intercept | 0.1655 | 0.8544 |
| Number of fruits | slope | 1.0699 | 0.8634 |
| Number of seeds per fruit | - | 0.0001 | 0.8527 |
| P(Recruitment) | - | 360.1242 | 4.4539 |
| Distribution of offspring size | mean | 0.1220 | 0.8539 |
| Distribution of offspring size | SD | 0.1135 | 0.8538 |

#### Appendix S3 Supplementary figures

**Figure S1.**
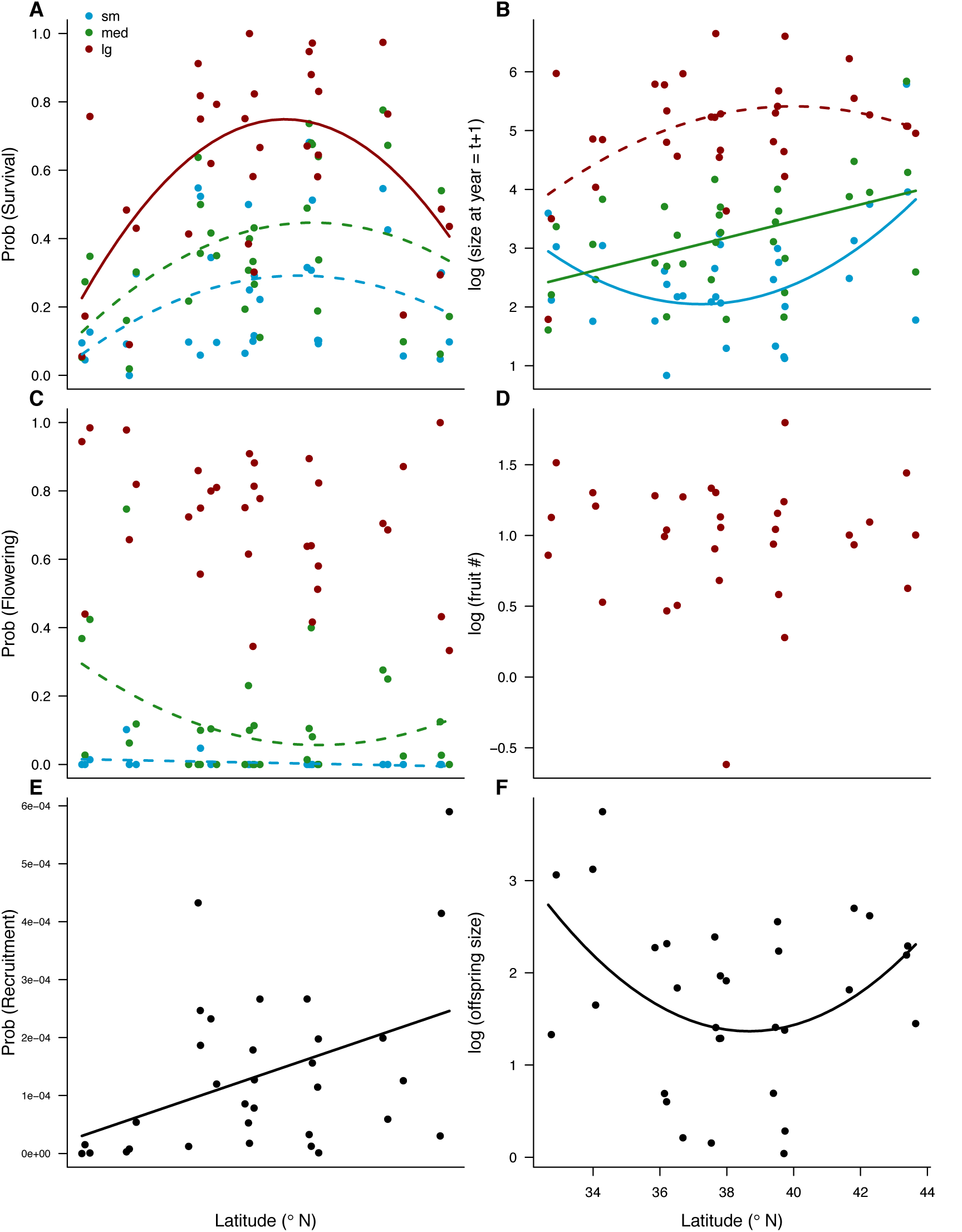
Vital rates as a function of latitude and elevation. Survival, growth, flowering, and fruit number are shown for small (within 0-20% quantile), medium (within 40-60% quantile), and large (within 80-100% quantile) plants. Solid lines represent slopes that are greater than 0 at P < 0.05, and dashed lines represent slopes that are greater than 0 at P < 0.1 (Table S4).

**Figure S2.**
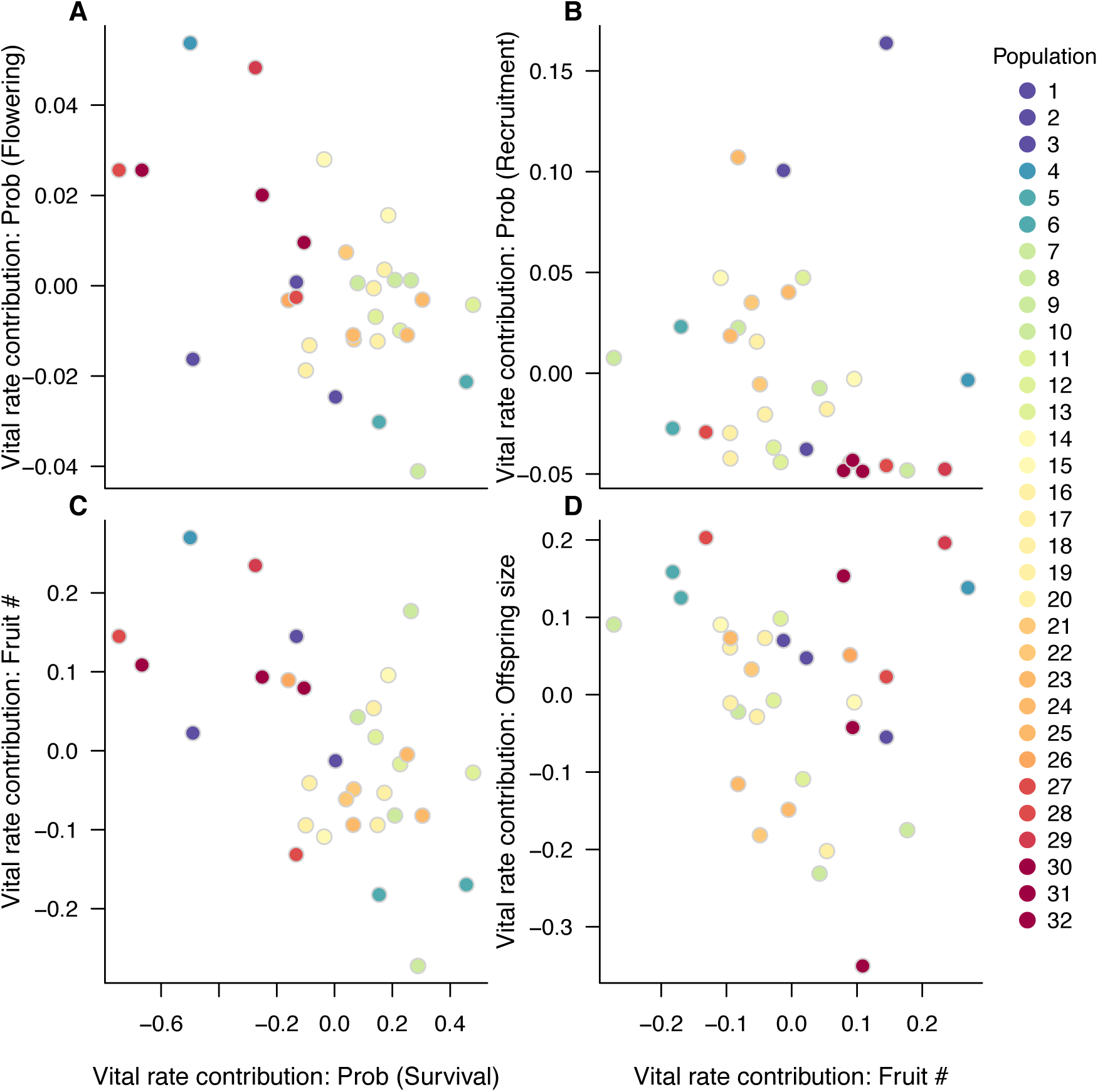
Population-specific vital rate contributions with significantly negative correlations *(P <* 0.05).

## REFERENCES

1. Sexton JP, McIntyre PJ, Angert AL, Rice KJ (2009) Evolution and ecology of species range limits. Annu Rev Ecol Evol Syst 40(1):415–436.

2. Brown JH (1984) On the relationship between abundance and distribution of species. Am Nat 124(2):255–279.

3. Sagarin RD, Gaines SD (2002) The “abundant centre” distribution: to what extent is it a biogeographical rule? Ecol Lett 5(1):137–147.

4. Abeli T, Gentili R, Mondoni A, Orsenigo S, Rossi G (2014) Effects of marginality on plant population performance. J Biogeogr 41(2):239–249.

5. Pironon S, et al. (2016) Geographic variation in genetic and demographic performance: new insights from an old biogeographical paradigm. Biol Rev. doi:10.1111/brv.12313.

6. Pironon S, Villellas J, Morris WF, Doak DF, Garcia MB (2015) Do geographic, climatic or historical ranges differentiate the performance of central versus peripheral populations? Glob Ecol Biogeogr 24(6):611–620.

7. Aikens ML, Roach DA (2014) Population dynamics in central and edge populations of a narrowly endemic plant. Ecology 95(7):1850–1860.

8. Villellas J, Doak DF, Garcia MB, Morris WF (2015) Demographic compensation among populations: what is it, how does it arise and what are its implications? Ecol Lett 18(11):1139–1152.

9. Doak DF, Morris WF (2010) Demographic compensation and tipping points in climate-induced range shifts. Nature 467(7318):959–962.

10. Read AF, Harvey PH (1989) Life history differences among the eutherian radiations. J Zool 219(2):329–353.

11. Charnov EL, Schaffer WM (1973) Life-history consequences of natural selection: Cole's result revisited. 791–793.

12. Cole LC (1954) The population consequences of life history phenomena. Q Rev Biol 29(2):103–137.

13. MacArthur RH, Wilson EO (1967) The theory of island biogeography (Princeton University Press, Princeton, NJ).

14. Southwood TRE (1977) Habitat, the templet for ecological strategies? J Anim Ecol 46(2):336.

15. Reinartz JA (1984) Life history variation of common mullein (*Verbascum thapsus*): I. Latitudinal differences in population dynamics and timing of reproduction. J Ecol 72(3):897.

16. Lacey EP (1988) Latitudinal variation in reproductive timing of a short-lived monocarp, *Daucus carota* (Apiaceae). Ecology 69(1):220–232.

17. Hewitt GM (1996) Some genetic consequences of ice ages, and their role, in divergence and speciation. Biol J Linn Soc 58(3):247–276.

18. Davis MB, Shaw RG (2001) Range shifts and adaptive responses to Quaternary climate change. Science 292(5517):673–9.

19. Svenning JC, Normand S, Skov F (2008) Postglacial dispersal limitation of widespread forest plant species in nemoral Europe. Ecography 31(3):316–326.

20. Hille Ris Lambers J (2015) Extinction risks from climate change. Science 348(6234):501–502.

21. Hargreaves AL, Samis KE, Eckert CG (2014) Are species' range limits simply niche limits writ large? A review of transplant experiments beyond the range. Am Nat 183(2):157–173.

22. Clausen JC, Keck DD, Hiesey WM (1948) Experimental studies on the nature of species. III. Environment responses of climatic races of Achillea. (Carnegie Institution of Washington Publication 581, Washington, D. C.).

23. Geber MA, Eckhart VM (2005) Experimental studies of adaptation in *Clarkia xantiana.* II. Fitness variation across a supspecies border. Evolution 59(3):521–531.

24. Angert AL, Schemske DW (2005) The evolution of species' distributions: reciprocal transplants across the elevation ranges of *Mimulus cardinalis* and *M. lewisii*. Evolution 59(8):222–235.

25. Sexton JP, Dickman EE (2016) What can local and geographic population limits tell us about distributions? Am J Bot 103(1):129–39.

26. Vaupel A, Matthies D (2012) Abundance, reproduction, and seed predation of an alpine plant decrease from the center toward the range limit. Ecology 93(10):2253–2262.

27. Jump AS, Woodward FI (2003) Seed production and population density decline approaching the range-edge of *Cirsium* species. New Phytol 160(2):349–358.

28. Eckhart VM, et al. (2011) The geography of demography: Long-term demographic studies and species distribution models reveal a species border limited by adaptation. Am Nat 178(S1):S26–S43.

29. Stanton-Geddes J, Tiffin P, Shaw RG (2012) Role of climate and competitors in limiting fitness across range edges of an annual plant. Ecology 93(7):1604–1613.

30. Lenoir J, Svenning J-C (2015) Climate-related range shifts - a global multidimensional synthesis and new research directions. Ecography 38(1):15–28.

31. Breshears DD, Huxman TE, Adams HD, Zou CB, Davison JE (2008) Vegetation synchronously leans upslope as climate warms. Proc Natl Acad Sci U S A 105(33):11591–2.

32. Svenning J-C, Sandel B (2013) Disequilibrium vegetation dynamics under future climate change. Am J Bot 100(7):1266–86.

33. Griffin D, Anchukaitis KJ (2014) How unusual is the 2012-2014 California drought? Geophys Res Lett 41(24):9017–9023.

34. Robeson SM (2015) Revisiting the recent California drought as an extreme value. Geophys Res Lett 42(16):6771–6779.

35. Diffenbaugh NS, Swain DL, Touma D (2015) Anthropogenic warming has increased drought risk in California. Proc Natl Acad Sci U S A 112(13):3931–6.

36. Angert AL (2006) Demography of central and marginal populations of monkeyflowers *(Mimulus cardinalis* and *M. lewisii*). Ecology 87(8):2014–2025.

37. Angert AL (2009) The niche, limits to species' distributions, and spatiotemporal variation in demography across the elevation ranges of two monkeyflowers. Proc Natl Acad Sci 106(Supplement 2):19693–19698.

38. Bayly M (2016) Translocations of *Mimulus cardinalis* beyond the northern range limit show that dispersal limitation can invalidate ecological niche models. doi:10.14288/1.0223357.

39. Samis KE, Eckert CG (2009) Ecological correlates of fitness across the northern geographic range limit of a Pacific Coast dune plant. Ecology 90(11):3051–3061.

40. Samis KE, López-Villalobos A, Eckert CG (2016) Strong genetic differentiation but not local adaptation toward the range limit of a coastal dune plant. Evolution 70(11):2520–2536.

41. Muir CD, Angert AL (2016) Grow with the flow: a latitudinal cline in physiology is associated with more variable precipitation in *Mimulus cardinalis*. bioRxiv. doi:10.1101/080952.

42. Leggett WC, Carscadden JE (1978) Latitudinal variation in reproductive characteristics of American shad *(Alosa sapidissima):* Evidence for population specific life history strategies in fish. J Fish Res Board Canada 35(11):1469–1478.

43. Hardie DC, Hutchings JA (2010) Evolutionary ecology at the extremes of species' ranges. Environ Rev 18(NA):1–20.

44. Kooyers NJ, Greenlee AB, Colicchio JM, Oh M, Blackman BK (2015) Replicate altitudinal clines reveal that evolutionary flexibility underlies adaptation to drought stress in annual *Mimulus guttatus*. New Phytol 206(1):152–165.

45. Olsson K, Ågren J (2002) Latitudinal population differentiation in phenology, life history and flower morphology in the perennial herb *Lythrum salicaria*. J Evol Biol 15(6):983–996.

46. Stearns SC (1983) The influence of size and phylogeny on patterns of covariation among life-history traits in the mammals. Oikos 41(2):173.

47. Salguero-Gómez R, et al. (2016) Fast-slow continuum and reproductive strategies structure plant life-history variation worldwide. Proc Natl Acad Sci U S A 113(1):230–5.

48. Salguero-Gómez R (2016) Applications of the fast-slow continuum and reproductive strategy framework of plant life histories. New Phytol. doi:10.1111/nph.14289.

49. Lee-Yaw JA, et al. (2016) A synthesis of transplant experiments and ecological niche models suggests that range limits are often niche limits. Ecol Lett 19(6):710–722.

50. Parmesan C, Yohe G (2003) A globally coherent fingerprint of climate change impacts across natural systems. Nature 421(6918):37–42.

51. Hickling R, et al. (2006) The distributions of a wide range of taxonomic groups are expanding polewards. Glob Chang Biol 12(3):450–455.

52. Baldwin BG, Goldman DH, Keil DJ, Patterson R, Rosatti TJ (2012) The Digital Jepson Manual: Vascular Plants of California, Thoroughly Revised and Expanded (Univ of California Press).

53. Easterling MR, Ellner SP, Dixon PM (2000) Size-specific sensitivity: Applying a new structured population model. Ecology 81(3):694–708.

54. Ellner SP, Rees M (2006) Integral projection models for species with complex demography. Am Nat 167(3):410–428.

55. Bates D, Mächler M, Bolker B, Walker S (2015) Fitting linear mixed-effects models using lme4. J Stat Softw 67(1):1–48.

56. R Core Team (2016) R: a language and environment for statistical computing. R Foundation for Statistical Computing, Vienna, Austria. Available at: http://www.r-project.org.

57. Williams JL, Miller TEX, Ellner SP (2012) Avoiding unintentional eviction from integral projection models. Ecology 93(9):2008–2014.

58. Merow C, et al. (2014) Advancing population ecology with integral projection models: a practical guide. Methods Ecol Evol 5(2):99–110.

59. Rees M, Childs DZ, Ellner SP (2014) Building integral projection models: a user's guide. J Anim Ecol 83(3):528–545.

60. Caswell H (2001) Matrix Population Models: Construction, Analysis, and Interpretation (Sinauer Associates, Sunderland, MA). 2nd Ed.

61. Wood SN (2011) Fast stable restricted maximum likelihood and marginal likelihood estimation of semiparametric generalized linear models. J R Stat Soc Ser B (Statistical Methodol 73(1):3–36.

62. Rees M, Ellner SP (2009) Integral projection models for populations in temporally varying environments. Ecol Monogr 79(4):575–594.

63. Ellner SP, Childs DZ, Rees M (2016) Data-driven modelling of structured populations: a practical guide to the integral projection model (Springer).

64. Wang T, Hamann A, Spittlehouse DL, Murdock TQ (2012) ClimateWNA - High-resolution spatial climate data for western North America. J Appl Meteorol Climatol 51:16–29.

65. Fackler PL (1991) Modeling interdependence: An approach to simulation and elicitation. Am J Agric Econ 73(4):1091–1097.

